# High contribution of canopy to oleoresin accumulation in loblolly pine trees

**DOI:** 10.1101/2020.04.09.034264

**Authors:** Amber N. Parrish, Glenn W. Turner, B. Markus Lange

## Abstract

The shoot system of all loblolly pine (*Pinus taeda* L.) contains abundant resin ducts, and the oleoresins contained within them have demonstrated roles in constitutive defenses. This study is providing a quantitative assessment of oleoresin biosynthesis and accumulation in resin ducts. Morphometric analyses of representative tissue sections indicate that the fractional volume of resin ducts is particularly high in the cortex of young stems and their needles, representing a major portion of total pine resins from primary growth of the canopy. We demonstrate that it is possible to extrapolate oleoresin formation from the microscopic scale (tissues sections) to the macroscale (entire trees), which has implications for assessing resins as renewable feedstocks for bioproducts.

## INTRODUCTION

According to recent assessments commissioned by the United States Department of Energy, roughly 320 million dry tons of biomass per year are currently removed sustainably from forestlands (Downing et al., 2011). As markets for bioenergy feedstocks are evolving, there is considerable potential for harvesting additional biomass from non-reserve forestland by removing logging residues and by strategic thinning. Woody biomass can contain high quantities of oleoresins, particularly in conifer forests. The chemistry of conifer oleoresins is characterized by mixtures of volatiles (primarily monoterpenes) and semi-volatiles (primarily diterpenoids) (Joye and Lawrence, 1967; Norin, 1972; Turner et al., 2019). These secretions are characterized by a high volumetric energy density and high degree of reduction. Because of these physicochemical properties, higher-boiling-point terpenoid resins are viable biocrude feedstocks for heavier liquid transportation fuels comparable to diesel and kerosene (Jiang and Zhang, 2018), while lower-boiling-point terpenoid oils are excellent feedstocks for the manufacturing of solvents and fine chemicals (Rubulotta and Quadrelli, 2019).

Oleoresins are part of both constitutive and inducible defense mechanisms in members of the pine family (Pinaceae) (Hudgins et al., 2004; Franceschi et al., 2005; Keeling et al., 2006). Pines produce oleoresin within a system of resin ducts that extends throughout the plant body. In the shoot, radial ducts extend from the vascular cambium across both the secondary phloem and secondary xylem (horizontally across the main stem), while the axial ducts extend along the stem axis (vertically along the main stem), each for a length of 10 cm or more (Werker and Fahn, 1969). Radial ducts are more numerous than axial ducts, but axial ducts tend to be larger, and are thus thought to contribute more significantly to resin production (Langenheim, 2003). The literature contains excellent descriptive work that provides a general depiction of the duct system of pine (Langenheim, 2003; Wu and Hu, 1997). However, none of these studies has provided quantitative estimates of resin duct volumes, and the present study was thus aimed at developing methods to predict oleoresin production by extrapolating from representative microscopic measurements of resin duct size and abundance.

## MATERIALS AND METHODS

### Plant Growth and Harvest

Loblolly pine seedlings (*Pinus taeda* L.) were obtained from a wholesale nursery (to capture tree-to-tree variation, these were not clones). Trees were maintained in a greenhouse under ambient lighting, with supplemental heating to 15°C on cold days (no other direct temperature control), and were approximately four years of age at harvest.

### Tissue Preparation for Microscopy

Freehand sections of short shoots and sliding microtome sections of larger long shoots provided the bulk of the observations, as described below. For resin embedded sections of developing short shoots, the shoots were dissected and treated overnight with a fixative containing 4 % (v/v) glutaraldehyde in 50 mM PIPES buffer (pH 7.2). Samples were post-fixed in aqueous 0.5 % (w/v) OsO_4_ for 2 h at 20°C and then dehydrated in an ethanol series. The ethanol was then exchanged for propylene oxide, with subsequent infiltration of Spurr’s resin (Ted Pella, Redding, CA, USA). Thick sections (1 µm) of long shoots were cut with glass knives using an Ultracut R microtome (Leica Microsystems, Buffalo Grove, IL, USA) and stained with toluidine blue. Sections were then photographed with an BH-2 Light Microscope (Olympus America Inc., Center Valley, PA, USA).

### Morphometric Analysis – Defining Flushes of Growth

Growth spurts of trees during spring in temperate climates are often referred to as a flushes of growth (FOGs). The greenhouse-grown loblolly pine trees employed in this study produced several flushes of growth per year. FOGs were defined by age, with FOG 1 signifying the newest growth, which was defined as the biomass between the tips of branches and the first whorl (or youngest stem segment) (Figure 1A). FOG2 was defined as the biomass between the first whorl and the second whorl. The same labeling system was employed for branches (with the newest growth again being defined as FOG 1), but additional information about where the branch originates (in Figure 1A, there is a branch at FOG 10) was retained in the numbering (Figure 1B). In addition, if further branching occurred (in Figure 1A, there is a branch at FOG 10, and further branching at FOG 3), that information was also retained in the numbering system (Figure 1B).

**Figure 1.**
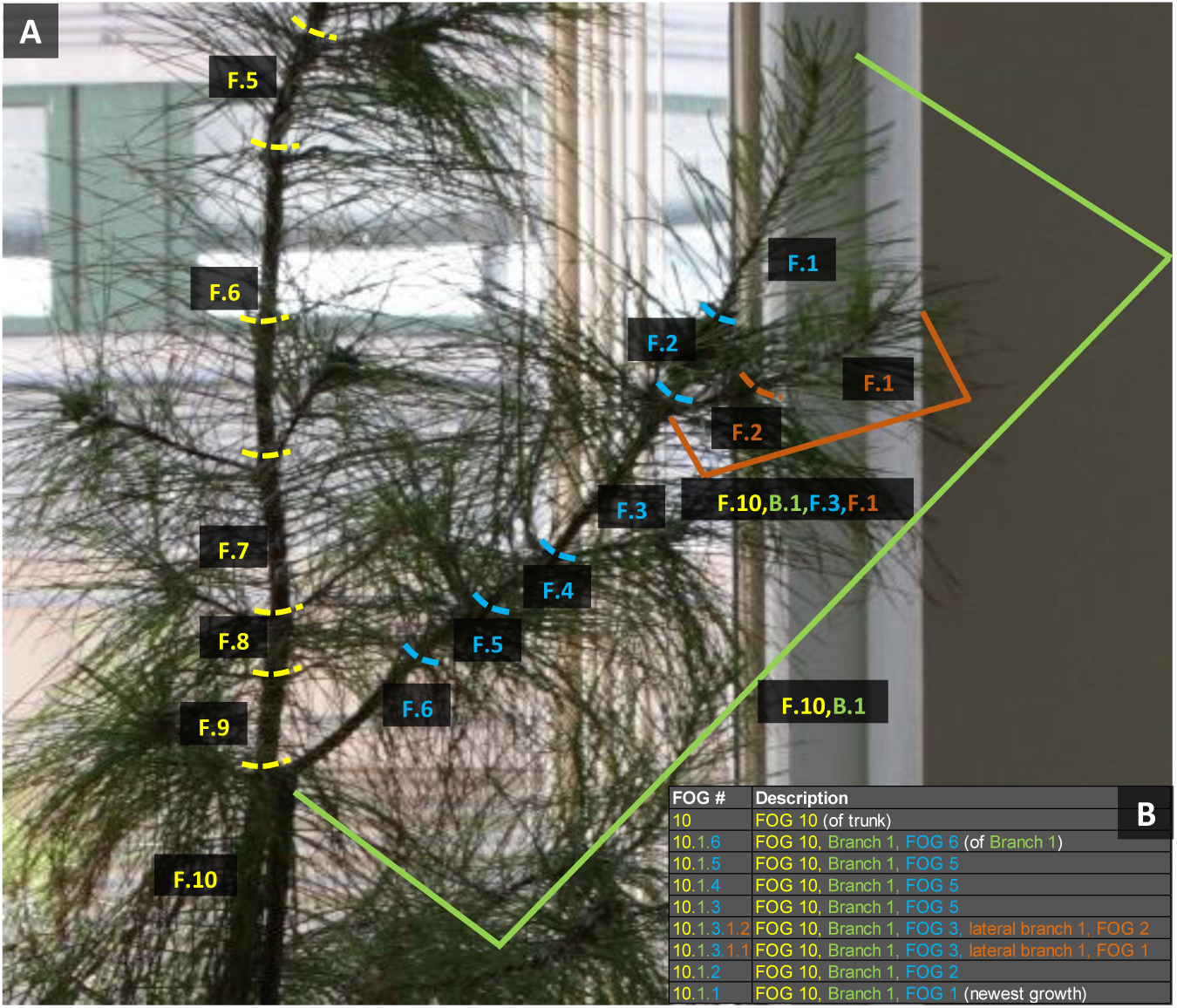
Definition of flushes of growth (FOGs). **A**, Pictorial description. **B**, Numbering system.

### Morphometric Analysis – Needle Measurements

Fascicles (bundles with a sheath typically holding 3 needles) were removed from the main stem and all branches. The number of needles was counted and their length recorded. The total needle biomass (fresh weight) was recorded as well. For each flush of growth, three representative needles, each from different fascicles, were picked at random. Using a razor blade,’ cross sections were obtained by hand in the center of needles. Sections were mounted on glass slides, a drop of mineral oil was added, and sections were viewed under a light microscope (DM LM, Leica Microsystems, Wetzlar, Germany) with camera attachment (Canon EOS Rebel XS, Tokyo, Japan). Sections were photographed with appropriate rulers for scaling. The area of the cross section was determined using the “Freehand” tool of (ImageJ, v.1.45s; https://imagej.nih.gov/ij/index.html; National Institutes of Health, USA) (Figure 2A). To convert the needle cross-sectional area into a volume, the tapering of needles at the distal end had to be taken into account (Figure 2B). The volume of the proximal two thirds was calculated by multiplying the cross-sectional area by the length of the needle up to that point (V = Area x Length). The volume of the remaining third was calculated using the formula for elliptical structures (V = ⅖ x Area x Length) (Figure 2C). The total volume of needles was then obtained by extrapolating the volume of an individual needle to that of all needles on each tree.

**Figure 2.**
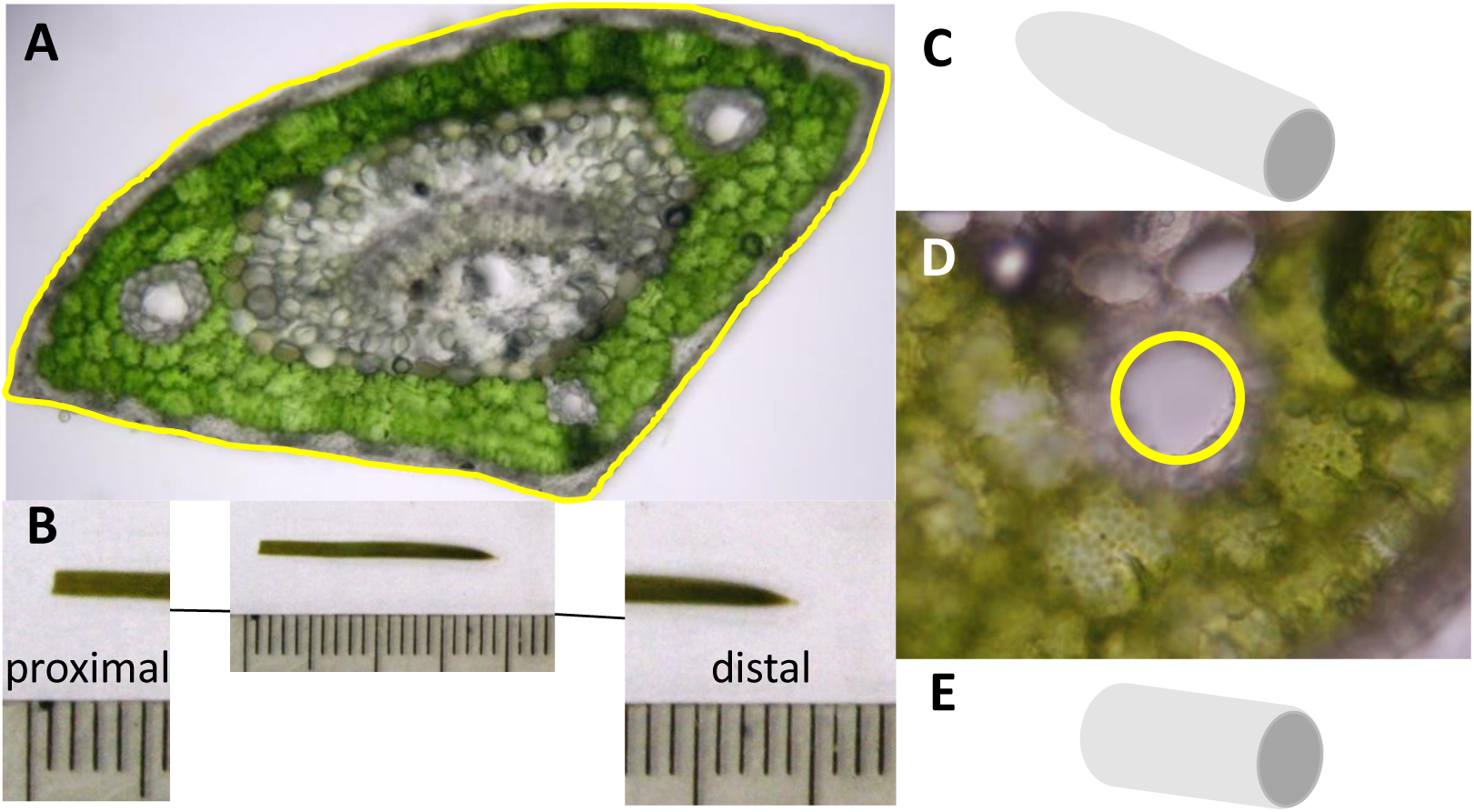
Morphometric analysis of needle features. **A**, Use of the “Freehand” function of ImageJ to obtain cross-sectional areas of needles. **B**, Tapering of needles at the distal end. **C**, Modeling the shape of the needles tip as elliptical cone. **D**, Use of the “Oval” function of ImageJ to obtain areas of resin ducts. **E**, Modeling the shape of a resin duct as cylinder.

The area of each resin duct was were determined using the “Oval” tool in ImageJ (Figure 2D). The resin duct volume was calculated by treating these structures as a cylinder that extends throughout the length of a needle (V = Area x Length) (Figure 2E). The total resin duct volume was estimated by calculating the volume of all resin ducts within a needle and then extrapolating to all needles on each tree.

### Morphometric Analysis –Stem Measurements

The length and circumference of the main stem and all branches were also recorded. Cross-sections, representing each FOG, were prepared by hand (with a razor blade) for smaller stems. Sections were then mounted on microscope slides with mineral oil to reduce shrinking/swelling of duct cells. The larger stems were sectioned cross-wise and tangentially using a sliding microtome and also mounted with mineral oil. Cross sections were performed by hand in the middle of a specimen (represented each FOG), sections placed on glass slides, and digital images taken as described above. Resin ducts occur primarily in the cortex and xylem of stems and a description of volume measurements is provided below.

#### Stem cortex

A hollow cylinder 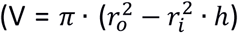 was assumed as a shape representing the cortex area of stems (Figure 3A). The relevant measurements were made using microscopic images using the “Straight Line” selection tool of ImageJ and are indicated in Figure 3B. For larger specimens, the number of cortical resin ducts was counted within the field of vision and then extrapolated to represent the number of cortical resin ducts per stem cross-section.

**Figure 3.**
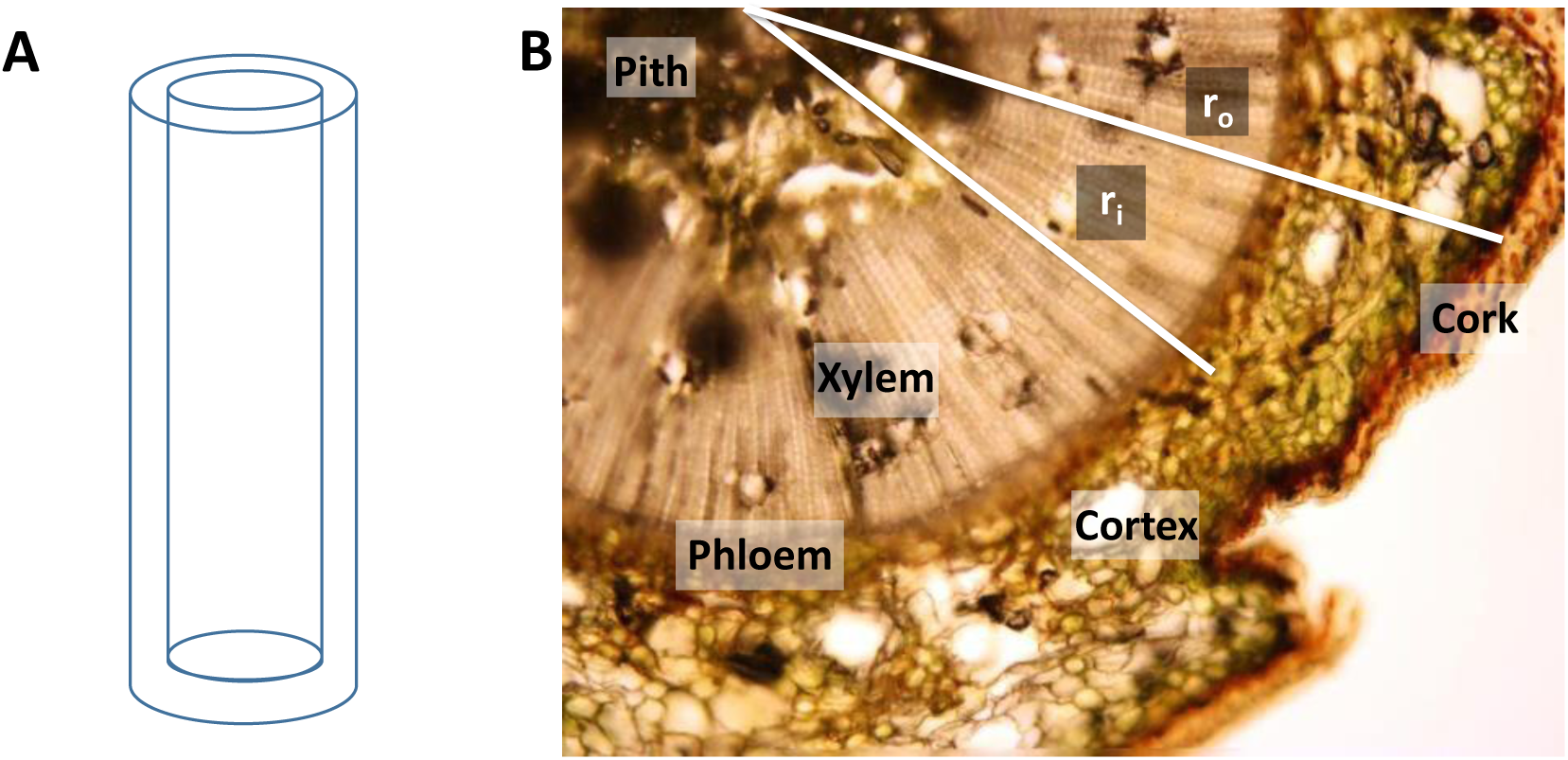
Estimation of stem cortex area. **A**, Shape of a hollow cylinder. **B**, Central pith-to-outer cortex radius (r_o_) and central pith-to-inner cortex radius (r_i_), which were used to determine the area of an annulus, which itself was used to calculate the cortex volume (multiplication with length of stem) as a hollow cylinder.

#### Stem xylem

Xylem tissue areas were estimated by tracing one-fourth of the xylem within a stem cross section using the “Freehand” function of ImageJ (Figure 4) and then multiplying the value by the factor four to obtain the entire area.

**Figure 4.**
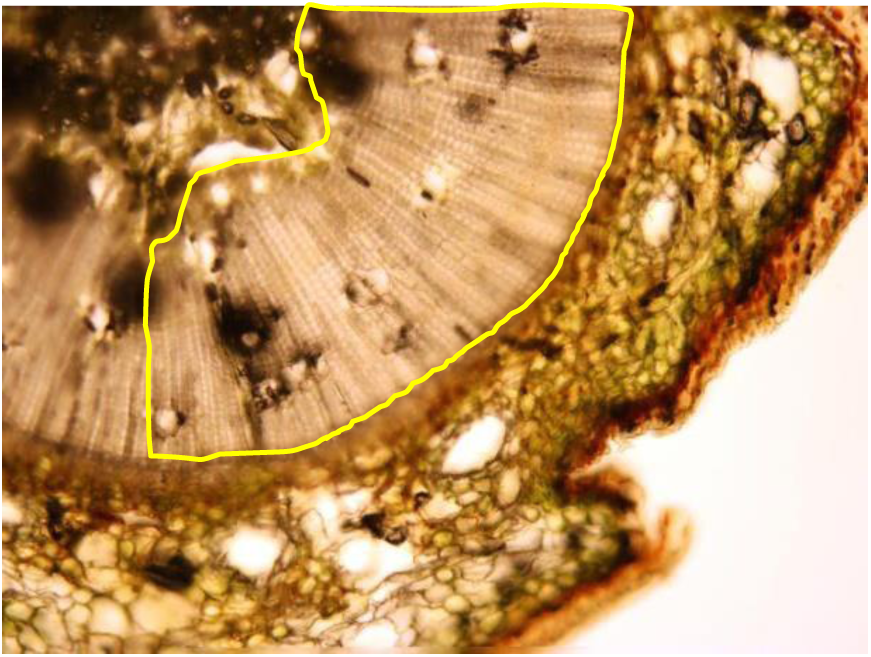
Estimation of stem xylem area. Highlighted in yellow is one-fourth of the xylem area, which excludes pith and phloem. The volume was determined by multiplication with the length of the stem.

### Morphometric Analysis – Xylem and Cortex Measurements

Cross sections were employed to visualize axial resin ducts (Figure 5A), while tangential sections served to capture radial resin ducts (Figure 5B). Numbers of ducts per section were tallied and extrapolated to obtain estimates for entire tissue sections, organs and trees. The “Oval” selection tool of ImageJ was used to calculate areas of axial and radial ducts (Figure 5C, D), with volumes being determined assuming a cylinder as the three-dimensional duct shape.

**Figure 5.**
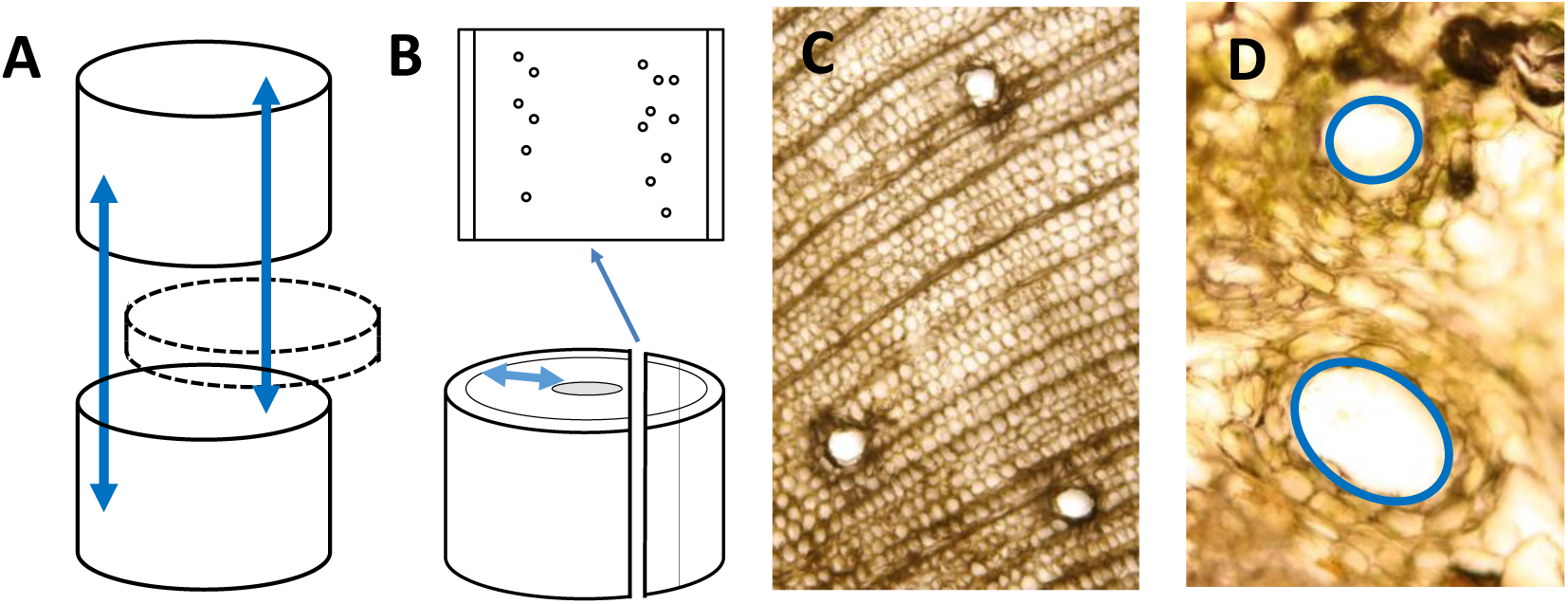
Estimation of axial and radial resin duct areas. **A**, Cross section to determine axial ducts (direction of ducts indicated by double-headed arrows). **B**, Tangential section to determine radial ducts (direction of ducts indicated by double-headed arrows). **C, D**, Use of the “Oval” function of ImageJ to obtain areas of resin ducts.

### Preparation of Samples for Oleoresin Analysis

**Xylem** and **cortex** were the primary tissues that contained resin ducts. While it was impractical to collect xylem and cortex separately for metabolite analyses, we noticed that cross sections of stems tended to rupture at the boarder of xylem and phloem tissues, which essentially left us with two samples types: wood (containing pith and xylem) and bark (containing phloem, cortex and cork), which were highly enriched in xylem and cortex, respectively. These tissue samples were processed separately to determine if differences in the chemical composition might exist. To obtain concentrations of metabolites, we determined the volumes of wood and bark as outlined below. The **wood** volume was determined using the “Straight Line” selection tool in ImageJ (to determine the relevant radii for an area estimation) and then multiplying by the length of the stem of the FOG of interest (assuming a cylinder as three-dimensional shape. A hollow cylinder 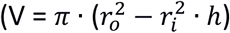 was assumed as a shape representing the **bark** area (see Figure 3A), where the radii are defined as *r*_*0*_ = radius from central pith to outer cork and *r*_*i*_ = radius from central pith to outer xylem (Figure 6).

**Figure 6.**
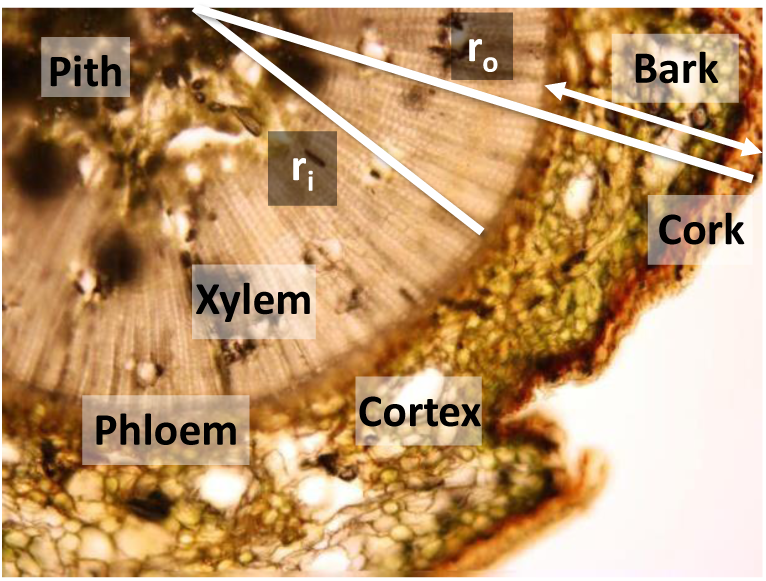
Estimation of wood area based on measuring relevant radii (white) using the “Straight Line” selection tool in ImageJ. For details see text.

## RESULTS

### High Resin Duct Volume in Needles and Cortex of Young Shoots

As a first step toward a detailed description of the loblolly pine resin duct system, we determined the biomass and volume of resin ducts in different parts of the plant. Briefly, four-year-old trees were harvested and separated into (i) main stem, (ii) short shoots on main stem, (iii) branches, and (iv) short shoots on branches. The main stem and branches were further dissected into woody parts (consisting mostly of secondary xylem) and the outer stem consisting of cortex or bark (mostly secondary phloem and cork layers). All tree parts were weighed and processed separately. Resin duct areas were estimated by quantitative image analysis and volumes calculated based on the physical dimensions of the original sample. The total resin duct volume was then estimated by extrapolation. The harvest (fresh) weights for the entire trees were 1,141 g (tree 1), 2,010 g (tree 2), and 2,344 g (tree 3) (Figure 7A). Averaged over three trees, the woody parts (combined from main stem and branches) and needles contributed about equally to biomass, while bark tissue of the main stem and branches was lighter (Figure 7B).

**Figure 7.**
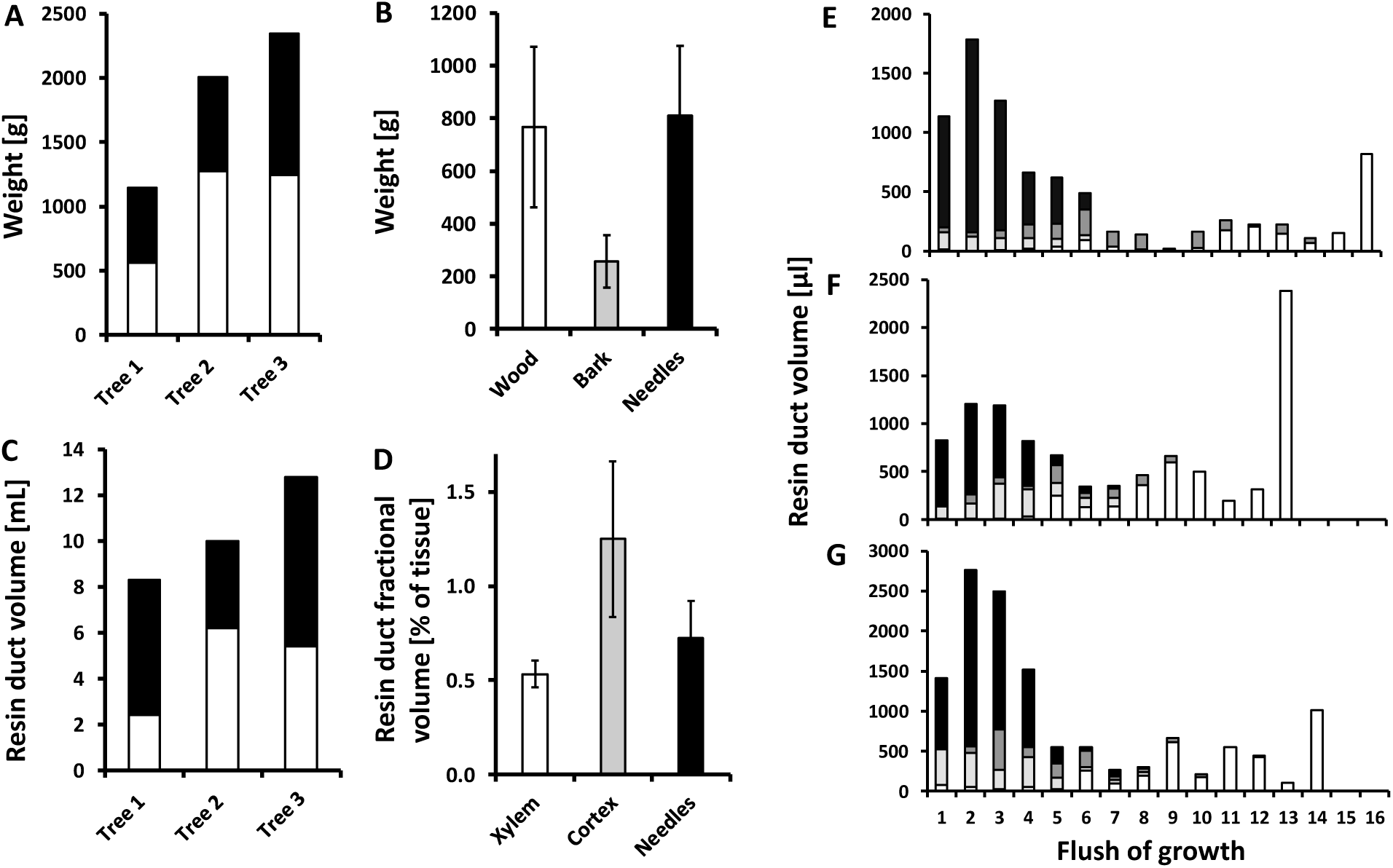
Biomass and Resin Duct Characteristics of Loblolly Pine Trees. Weight of main stem plus branches (white bars) and needles (black bars) of individual trees (**A**). Weight of different tree parts, averaged across trees (n = 48; bars, standard deviation) (**B**). Resin duct volume of main stem plus branches (white bars) and needles (black bars) of individual trees (**C**). Fractional volume of resin ducts in different tissues, averaged across trees (n = 48; bars, standard deviation) (**D**). Volume of resin ducts, separated by flushes of growth (**E-G**). Different parts of trees are indicated with grayscale bars (white, main stem; bright gray, branches; dark gray, needles attached to main stem; and black, needles attached to branches) (tree 1, **E**; tree 2, **F**; tree 3, **G**).

Based on morphometric analyses of representative microscopic images, the total resin duct volume was calculated, by extrapolation, to amount to 8.2 ml for tree 1, 9.9 ml for tree 2, and 12.8 ml for tree 3 (Figure 7C). Needles (from both main stem and branches) contained a very significant proportion of the resin duct volume in tree 1 and tree 3 (5.8 and 7.4 ml, respectively), while the main stem and branches had lower volumes (2.4 and 5.4 ml for tree 1 and tree 3, respectively). In tree 2, the main stem and branches were the largest contributors to resin duct volume (6.2 ml), with needles containing less (3.7 ml). We then calculated the contribution of resin ducts to the overall volume of each organ and tissue type. Resin ducts of the main stem and branches were primarily found in secondary xylem tissue, with a smaller contribution from the cortical layers of primary tissues near the apices of the trees. The fractional volume of resin ducts (as percentage of volume of tissue type) was highest in the cortex of the stem and branches (1.25 ± 0.96 %), followed by needles (0.76 ± 0.31 %), and then xylem of stem and branches (0.53 ± 0.15 %) (Figure 7D).

### Relative Contribution of Resin Duct Types Differs Across Flushes of Growth

To assess developmental patterns of resin accumulation, tree parts were separated by flushes of growth (FOGs; spurt of growth occurring periodically) and resin duct volumes calculated based on analyses of microscopic images. In tissue samples taken from the oldest FOGs (11-16), the resin duct volume was dominated by the contribution from the main stem (65 - 100 %) (Figure 7E-G). Needles directly connected to the main stem made highly variable contributions (0 – 81 %) to the total resin duct volume for all other FOGs (1-10). In the youngest FOGs (1-4), the relative volume of resin ducts of stems and needles on branches was particularly high (63 - 91 %) (Figure 7E-G). For FOGs 1-4, the combined resin duct volume of needles (on main stem and branches) accounted for 61-93 % of the total resin duct volume (in all trees).

### Architecture of the Resin Duct System

Typically, three needles occurred in fascicles on short shoots. Needles had a central vascular cylinder consisting of two vascular bundles surrounded by transfusion tissue, all enclosed by an endodermis. The vascular cylinder was surrounded by a photosynthetic mesophyll layer, situated between the endodermis and a sclerified hypodermal layer. In cross sections, the vascular cylinder appeared oval and moderately flattened dorsiventrally, extending laterally to about ten cell layers from the needle edge (Figure 8A-C).

**Figure 8.**
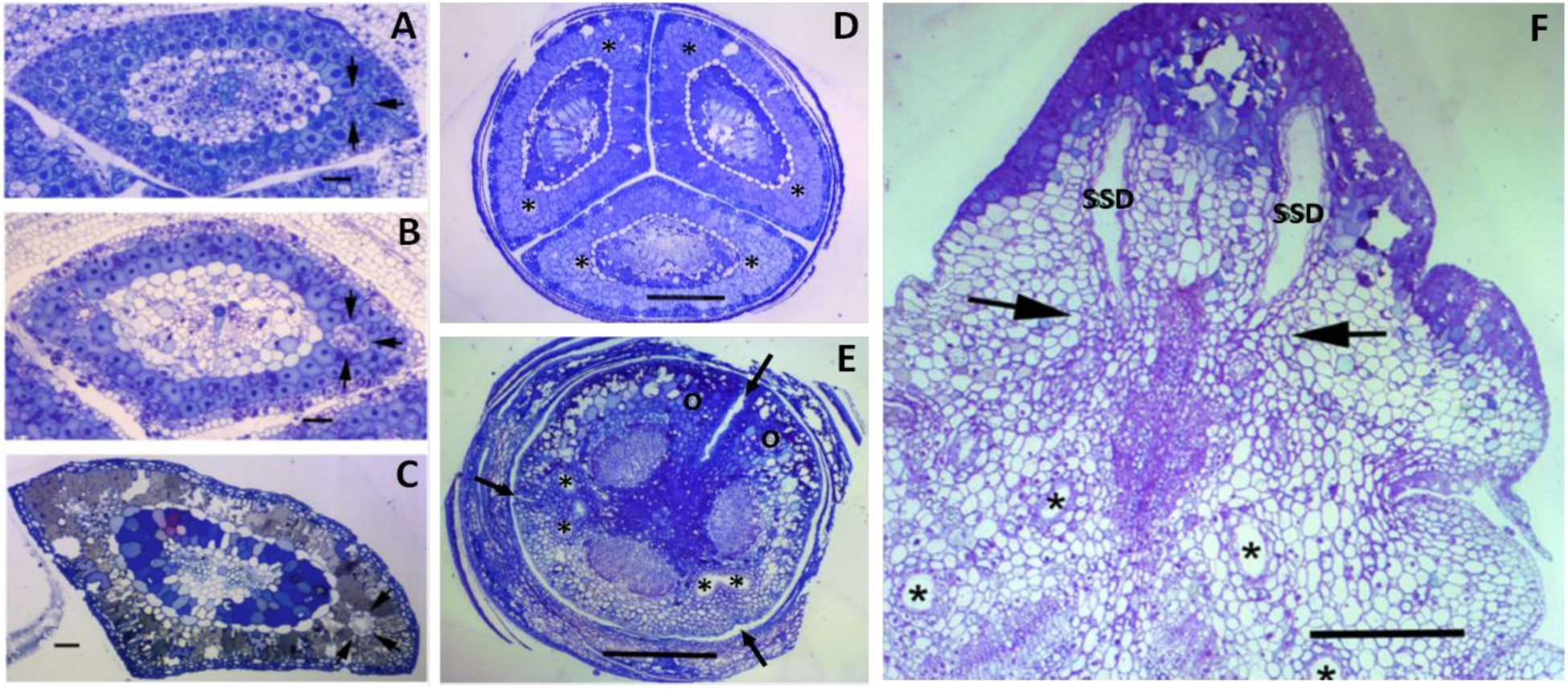
Architecture of the loblolly pine resin duct system. Transverse sections of loblolly pine needles representing a young portion near the intercalary meristem (**A**), an older portion of lateral resin ducts at early secretory phase (**B**), and a mature needle with post-secretory phase resin ducts (**C**). Arrows indicate one of the two lateral initiating resin ducts; bars = 100 µm. (**D**) Cross section of a developing fascicle showing three young needles still ensheathed by scale needles (bar = 500 µm). The positions of resin ducts are indicated by asterisks. (**E**) Cross section through a region of a developing fascicle where the needle bases connect to the short shoot and where branched cortical resin ducts of the short shoot (*) connect with lateral resin ducts of the needles (O) (bar = 500 µm). Arrows indicate clefts that mark the separation of different needles. (**F**) Cross section of a young long shoot branch at a juncture where a lateral short shoot attaches. A portion of the short shoot can be seen above the arrows. Cortical ducts of the long shoot are indicated with (*) and short shoot cortical ducts are marked with (SSD) (bar = 500 µm).

In our young, greenhouse-grown trees, two resin ducts occurred consistently in the mesophyll, one along each lateral side of the vascular cylinder and one cell layer removed from the endodermis. In some cases, needles had two additional resin ducts bordering the center of vascular cylinder, on both the abaxial and adaxial sides. During growth, new tissues were generated from intercalary meristems at the base of the needles, so that mature tissues were found the central and apical areas of the needle, while newly differentiated tissues were found near the needle base. Mature resin ducts formed long channels that extended into the needle base and connected directly to cortical resin ducts of the short shoots (Figure 8E).

The short shoots are determinate stems, each connecting a fascicle with the main stem. Each short shoot contained three cortical resin ducts, which bifurcated immediately below the base of the needles. The lumen of each cortical duct was found to be directly connected to a lateral resin duct of a different needle, so that each short-shoot cortical duct is connected to two different needles and the two lateral ducts of each needle connect to two different cortical ducts (Figure 8D, E). Cortical ducts appear to terminate at an abscission zone at the base of the short shoot and do not connect with the large cortical ducts of the main primary stems (Figure 8F). Therefore, the needle ducts can be considered a separate compartment from the large cortical ducts of the main stems, which was an important observation that guided our calculations of resin duct volumes.

### Oleoresin Estimations Based on Morphometric Measurements are in Good Agreement with Those Based on Chemical Analyses

The quantities of known chemical constituents of pine oleoresin – monoterpenes, sesquiterpenes diterpene and resin acids – were determined from approximately one hundred representative tissue samples by gas chromatography coupled with flame ionization detection (GC-FID) (detailed methods description in Turner et al., 2019). The oleoresin content, quantified by chemical analyses of the same tissue samples, was then extrapolated to entire trees by considering their physical dimensions, which allowed us to also estimate the contribution of needles (2.8, 2.9 and 5.2 ml for tree 1, 2 and 3, respectively), woody parts (1.8, 6.3 and 4.5 ml for tree 1, 2 and 3, respectively), and bark (0.8, 2.6 and 1.2 ml for tree 1, 2 and 3, respectively); these values were very similar to those extrapolated from morphometric analyses of microscopic images (Figure 9).

**Figure 9.**
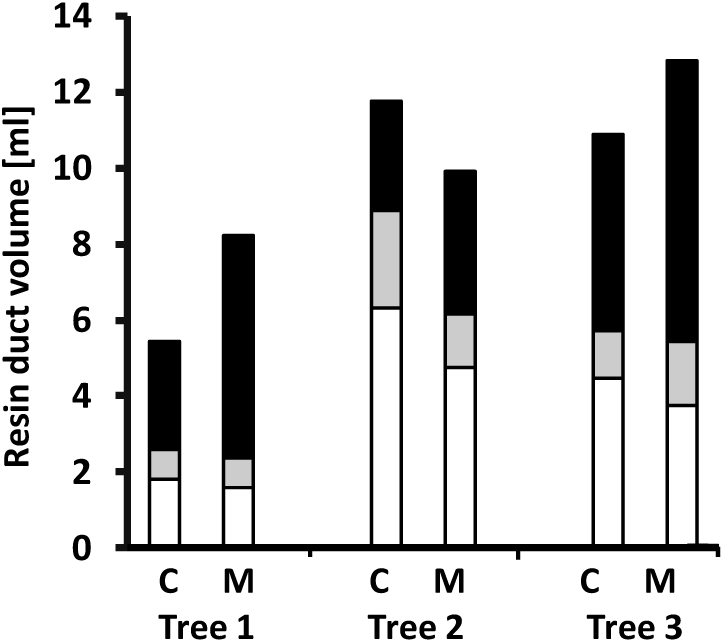
Estimation of resin content in loblolly pine trees. Comparison of oleoresin yields (in ml) as extrapolated from quantitative terpenoid measurements (C) and resin duct volume estimated based on microscopic analyses (M). Bars represent woody parts of the stem and branches (white), bark of the stem and branches (gray), and needles of the stem and branches (black).

We then asked if there were age-related differences in the accumulation of different classes of terpenoids (focusing on monoterpenes and diterpene resin acids; sesquiterpenes were not analyzed due to their low abundance). We divided trees up by growth segments, where segment 16 would correspond to the oldest, basal, part of each tree and segment 1 to the apex. Our measurements indicated that there was a negative, linear correlation between tissue age and diterpene resin acid content in needles and woody tissues (Figure 10). Samples that contained older tissues had lower diterpene resin acid concentrations, with a similar, but statistically not significant, trend in bark samples (Table 1). For monoterpene volatiles we observed a negative correlation between age and resin constituent in bark and needles, but not in xylem (Table 1).

**Figure 10.**
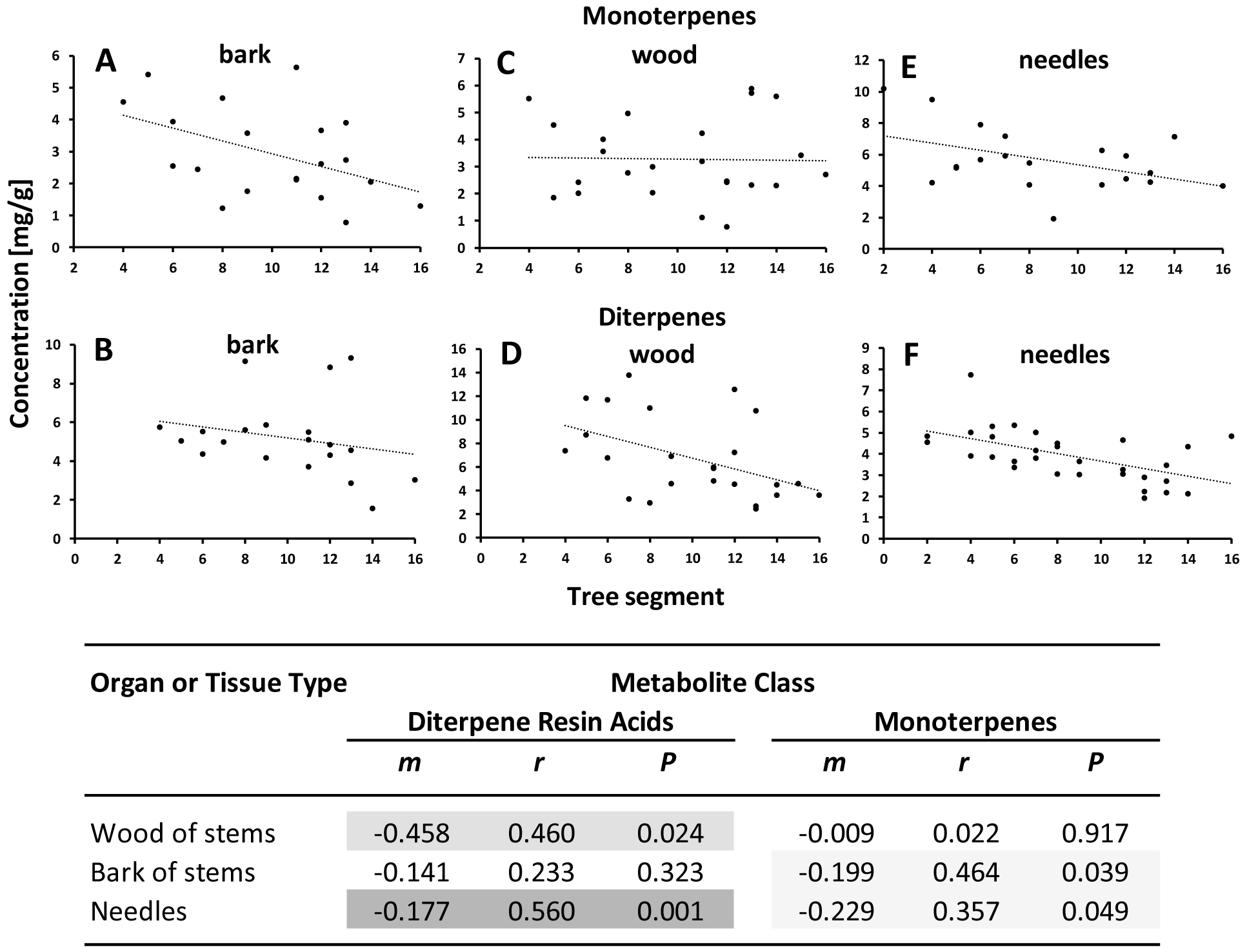
Developmental regulation of the accumulation of monoterpenes (panel **A, C** and **E**) and diterpene resin acids (panels **B, D** and **F**) in loblolly pine trees (segments are numbered 1-16, with 1 being the youngest and 16 the oldest growth material). The table at the bottom summarizes the correlation analyses between organ/tissue and oleoresin composition. Parameters of the statistical analysis are m, slope of correlation trendline; r, Pearson correlation coefficient; and P, Student’s t-test P-value. Gray background is used to indicate statistically significant correlations.

## DISCUSSION

### Developmental Gradient of Oleoresin Accumulation

We found that quantities of diterpene resin acids in young stems and needles are larger than the amounts contained in older shoots. Monoterpene volatile contents in bark of stems and needles, but not xylem of stems, were also larger in younger tissues (Table 1). These data indicate a more significant contribution of resins from primary tissues than from secondary tissues in four-year-old trees. Oleoresin accumulation has been demonstrated to be correlated with defense reactions and resistance to insect attack (Franceschi et al., 2005; Keeling and Bohlmann, 2006), but several studies have indicated that there is a trade-off between investment in growth and defenses in pine (Lorio and Sommers, 1986; Sampedro et al., 2011). It has been argued that the biosynthesis of reduced terpenoids comes at a high metabolic cost (Gershenzon, 1994), and it would seem that regulatory mechanisms have evolved that favor a proportionally higher investment in oleoresin deposition early during tissue development. This knowledge can help with identifying genes that are responsible for regulating oleoresin accumulation, which in turn can support the genomic selection of trees for improved resistance to bark beetle infestation and biofuel feedstocks (Westbrook et al., 2013, 2015).

### Canopy is a Major Contributor to Resin Accumulation in Loblolly Pine

The literature contains excellent qualitative descriptions of the resin duct system of pine stems and needles (Werker and Fahn, 1969; Fahn and Benayoun, 1976; Bosshard and Hug, 1980; DeAngelis et al., 1986; LaPasha and Wheeler, 1990; Blanche et al., 1992; Wu and Hu, 1997). However, this is the first study to quantitatively describe resin duct contents for the entire shoot system of young pine trees. Total resin duct volumes for specific organs and tissue types as well as entire trees were extrapolated from data obtained by two complementary approaches: (1) morphometric analyses of representative microscopic images and (2) oleoresin chemical analyses of more than one hundred tissue sections. The quantitative outcomes of these estimations were very similar. Both methods predicted a major contribution of needles (55.6 ± 16.8 % by morphometrics; 41.4 ± 14.8 % by chemical analysis) to the overall quantity of oleoresin. The majority of resin duct studies with pine species have focused on stems (Langenheim, 2003), as has the commercial exploitation of resins (Hillwig et al., 2010; da Silva Rodrigues-Corrêa et al., 2012). It had been suggested by Kelkar et al. (2006) that resin collection from smaller tress, branches and needles is a neglected, value-adding, source of revenue for commercial loblolly pine plantations. These logging residues, whose biomass has a higher contribution from needles than those of mature trees, are often burned, with only limited value as soil fertilizer. We are providing quantitative estimates of resin content in small loblolly pine trees, which reflect the properties of the canopy of larger trees (Naidu et al., 1998; da Silva Rodrigues-Corrêa et al., 2012). Our findings thus have implications for evaluating the potential of using forest biomass for biofuel and biomaterial applications.

## AUTHOR CONTRIBUTIONS

B.M.L. and G.W.T. conceived of the project. B.M.L., G.W.T., and A.N.P. designed experiments and analyzed data. A.N.P. and G.W.T. performed the experiments. B.M.L. wrote the manuscript, with input from the other authors. All authors read and approved the final manuscript.

## ACKNOWLEDGMENTS

This work was supported by the Division of Chemical Sciences, Geosciences, and Biosciences, Office of Basic Energy Sciences, and US Department of Energy (Grant No. DE-SC0001553 to B.M.L.). The authors would like to thank the Franceschi Microscopy & Imaging Center at Washington State University for access to instruments. No conflict of interest declared.

